# Resting-state fMRI foundation models enable robust and generalizable latent neural target discovery in cognitive aging interventions

**DOI:** 10.64898/2025.12.30.697042

**Authors:** Xinliang Zhou, Meishan Ai, Ehsan Adeli, Yu Zhang, Yang Merik Liu, Feng Vankee-Lin

## Abstract

The benefits of interventions targeting cognitive aging vary substantially across individuals, largely owing to heterogeneity in aging-related comorbidities. It is necessary to robustly identify neural patterns underlying intervention response and test their generalizability across heterogeneous cohorts. Resting-state functional MRI (rsfMRI) offers a potential pathway, but relying on predefined summary features with conventional methods has limited capacity to capture both within-individual longitudinal variation and between-individual differences, particularly in small and heterogeneous studies. Recent rsfMRI foundation models pretrained on large observational cohorts present a promising alternative by learning transferable spatiotemporal representations from time-series signals. Yet their validity and generalizability in local intervention settings remain unclear. Here, we systematically evaluated rsfMRI foundation models using data from two independent randomized controlled trials of older adults with mild cognitive impairment, testing whether these models can robustly extract longitudinal brain representations that predict post-intervention changes in episodic memory across trials. Foundation models outperformed conventional machine learning and deep learning approaches across both trials. Clinically informed adaptation using an external Alzheimer’s disease cohort further improved performance and robustness to confounders (i.e., head motion, site, and intervention arm), with accuracy up to 82%. Multivariate decomposition of foundation model embeddings identified latent neural patterns associated with episodic memory change with cross-study consistency at baseline that became more spatially distributed at post-intervention. These findings show that rsfMRI foundation models can enable robust and generalizable identification of latent neural patterns linking longitudinal brain dynamics to individual intervention response, laying the foundation for precision-driven neural target discovery in cognitive aging research.

**Significance Statement:** Interventions for cognitive aging show highly variable outcomes across individuals, limiting their clinical effectiveness. This study introduces a foundation model-based approach to identify latent neural patterns underlying individual differences in intervention response using resting-state fMRI. By leveraging pretrained models and domain-adaptive fine-tuning, we demonstrate robust and generalizable prediction of cognitive improvement across independent trials. Our findings suggest that latent brain representations, rather than predefined features, provide a scalable pathway toward precision-driven intervention strategies for aging populations.

## Introduction

As the global population ages, scalable interventions with minimal side effects, such as cognitive training and aerobic exercise, has become a focus in clinical research aimed at preventing cognitive decline^1^. However, the cognitive benefits of these interventions highly vary across individuals due to individual heterogeneity in aging-related comorbidities^2^. This variability has been a primary contributor to a history of inconclusive or failed trials^3–5^. However, rather than viewing these failures as mere setbacks, they represent a critical source of data. The accumulation of mixed results underscores the need to move beyond a trial-and-error approach. By systematically analyzing data across diverse trials, it may be possible to identify robust, generalizable patterns of neuroplastic potential, i.e., the brain capacity for beneficial cognitive change, paving the way for the future more consistently effective and individually tailored strategies for cognitive enhancement.

Resting-state functional magnetic resonance imaging (rsfMRI) provides an objective approach to examine individuals’ neuroplastic potential^6,7^. Traditionally, the utility of rsfMRI for mechanistic understanding of interventions has relied on the assumption that neuroplastic potential from interventions manifest within pre-defined regions of interest (ROIs) or at the level of pairwise functional connectivity (FC)^8^. This assumption imposes a restrictive representational framework. By aggregating BOLD signals into static ROI-level or connectivity summaries, such approaches may obscure complex, higher-order interactions among distributed neural components^9^, where intervention-related mechanisms may instead be expressed in latent, multivariate patterns rather than localized changes in individual ROIs or discrete FC edges^10^. This limitation is particularly consequential in aging cohorts, where intervention-related neural responses tend to be heterogeneous, dynamic, and idiosyncratic, diminishing the sensitivity of fixed anatomical targets or predefined connectivity metrics for capturing individual-level response signatures^2^. Analytically, many statistical and machine learning (ML) approaches to rsfMRI operate directly on these pre-defined features^11^. Conversely, conventional deep learning (DL) that operate on raw time-series data often require massive datasets that are often unavailable in real-world clinical settings^12^. In limited-data regimes, DL models may either oversimplify temporal dependencies or overfit to noisy BOLD signals, further constraining their utility for mechanistic inference^13^.

To uncover these understudied latent, multivariate neural patterns robustly across studies, we propose a new approach that leverages Foundation Models (FMs) to learn brain representations directly from raw rsfMRI timeseries. FMs, which are pre-trained on vast and diverse datasets, learn fundamental, generalizable patterns of brain activity before being applied to specific downstream tasks^14–16^. The strength of this approach lies in three key areas: First, FM architectures, often Transformer-based, are explicitly designed to capture complex, long-range temporal dependencies in BOLD signals that are lost in static FC summaries^17^. Second, the extensive pre-training provides a strong inductive prior, enabling these models to extract meaningful spatiotemporal dynamics for smaller, local intervention datasets while mitigating overfitting. By pre-training on thousands of hours of rsfMRI data, FMs can distill robust and generalizable neural representations that preserve fine-grained spatiotemporal signatures of brain function^18,19^. Third, FM-derived embeddings offer a compact, low-dimensional space that can be linked to behavioral outcomes, facilitating the discovery of latent neural variables. This capacity is particularly useful in intervention studies of aging, where neural responses are heterogeneous and individualized. However, a systematic methodology for adapting FMs to challenging small-sample settings, such as local, aging-focused intervention studies, has not yet been established. Therefore, the central goal of our work is to develop and validate such an approach, demonstrating how FMs can be reliably adapted to identify latent patterns of neuroplastic potential in older adults and support data-driven brain mechanism discovery.

In this paper, we aim to identify latent, multivariate neural patterns that are robust across intervention studies by adapting rsfMRI FMs and evaluating their utility in predicting individual cognitive responses in aging populations. Specifically, we address four research questions relevant to applying rsfMRI foundation models in local cognitive aging intervention studies: (Q1) whether FM-derived embeddings improve predictive performance over conventional approaches in small, heterogeneous cohorts; (Q2) whether clinically informed domain adaptation and joint modeling of baseline and post-intervention states enhance intervention-relevant representation; (Q3) whether FM-derived representations remain robust under common neuroimaging confounders including site variability, head motion, and intervention arm; and (Q4) whether their latent structure shows consistent organization across independent intervention cohorts.

To answer these questions, we evaluated two representative rsfMRI FMs: BrainLM^18^, trained to model whole-brain activity as an inference sequence similar to a language model, and BrainJEPA^19^, designed to learn shared embeddings from neighboring brain states. Both rsfMRI FMs were originally pre-trained on large-scale datasets, such as UK Biobank^20^ and the Human Connectome Project (HCP)^21^, which predominantly consist of younger, neurologically healthy adults. We further fine-tuned them using the Alzheimer’s Disease Neuroimaging Initiative (ADNI)^22^ dataset to ensure the brain representations reflect the range of variability covering both normal and abnormal cognitive aging. We then tested the adapted FMs on two independent local intervention studies targeting cognitive aging: ACT^23^ study (NCT03313895), a three-site randomized control trial (RCT) with four arms over six-month exercise and cognitive training combined, alone, or active control, and CogTE^24^ study (NCT02559063), a single-site RCT with two arms over six-week cognitive training or active control. Notably, both trials exemplify a common challenge in aging research: neither study identified a statistically significant group-level improvement in their primary outcome, episodic memory (EM) in older adults with mild cognitive impairment (MCI). This very absence of a clear average treatment effect makes them ideal testbeds for our purpose. The mixed results highlight the failure of traditional group-level analyses and create a need for methods that can uncover the individual-level neural factors distinguishing those who benefited from those who did not. To this end, we used rsfMRI data collected at baseline and post-intervention from both studies. Change in EM (ΔEM) was calculated with responders defined as sustained/improved post-intervention from baseline (ΔEM>=0) vs. non-responders as declined cases (ΔEM<0). By focusing on individual-level outcome, we explicitly test FM-derived brain representations can capture latent, multivariate neural patterns that robustly explain variability in cognitive response across studies. The overview of the analytical flow is presented in **Figure 1**.

**Figure 1.**
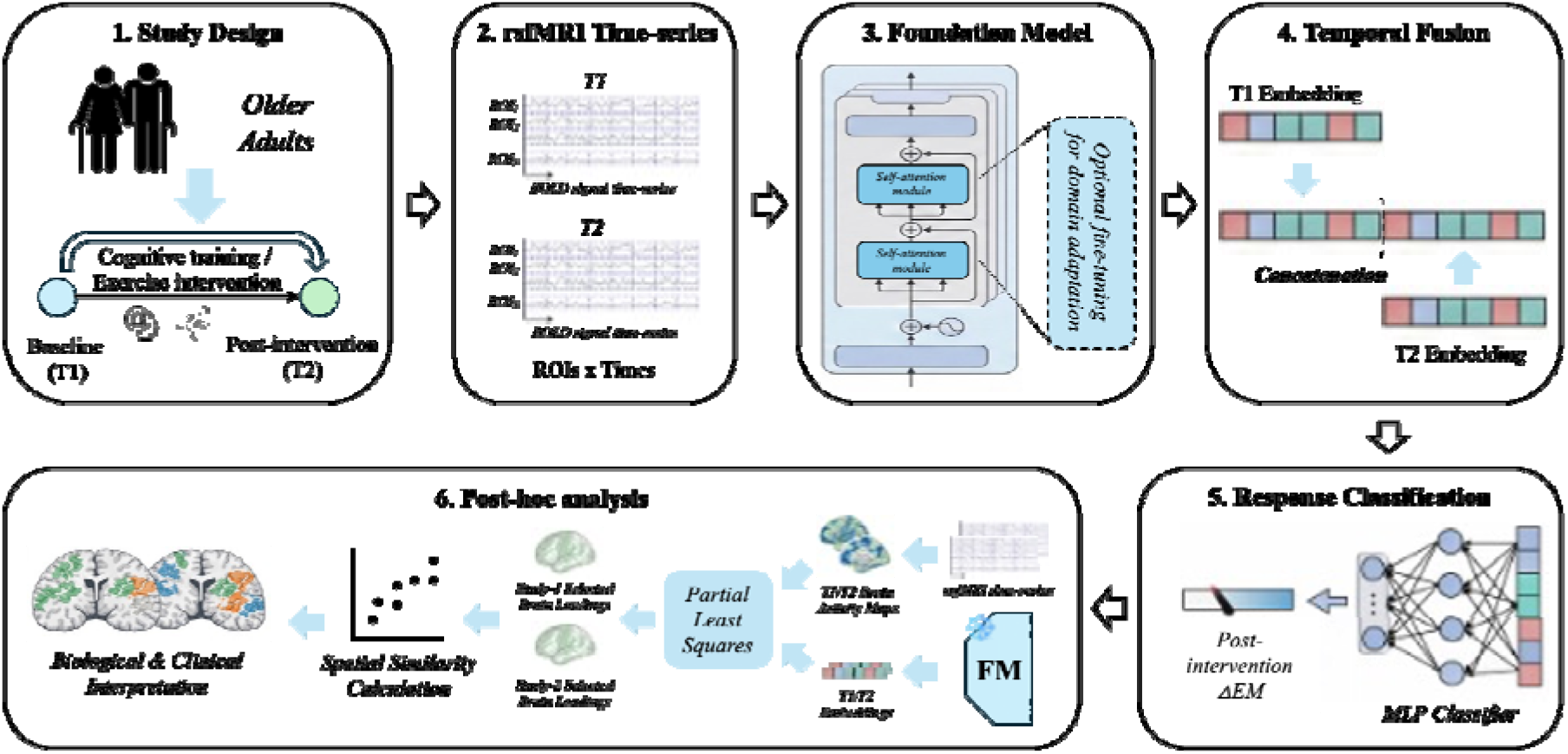
Schematic overview of the analytical pipeline. rsfMRI data collected at baseline (T1) and post-intervention (T2) were independently processed through a dual-stream architecture with shared weights. Pretrained FMs (BrainLM or BrainJEPA) operate directly on regional-averaged BOLD time-series, which are temporally segmented into patches to extract latent whole-brain representations at T1 and T2 respectively. The resulting T1 and T2 embeddings were concatenated to explicitly encode longitudinal neural change and used as input to a lightweight downstream classifier trained to infer EM. Following model training and evaluation within a stratified cross-validation framework, post-hoc multivariate analyses were performed to characterize latent embedding-brain associations and assess their consistency across independent studies. **NOTE:** rsfMRI, resting-state functional magnetic resonance imaging; FM, foundation model; BOLD, blood-oxygen-level–dependent; T1/T2, baseline/post-intervention timepoints; MLP, multilayer perceptron; EM, change of epsiodic memory; P, participant; RDM, representational dissimilarity matrix.

## Results

### Classification of change of episodic memory after intervention: Benchmark comparisons

To address Q1, we first evaluated whether rsfMRI foundation models can provide more informative brain representations than conventional approaches for predicting individual intervention response. Evaluation was performed under stratified five-fold cross-validation, balancing outcome labels and key study factors (intervention arm and site when applicable). The performance of two FMs, BrainLM and BrainJEPA, was systematically compared with three widely used data-driven approaches: Support Vector Machine (SVM)^25^, a ML method using statistical learning theory; Multilayer Perceptron (MLP)^26^, a DL method capturing nonlinear relationships; and Graph Convolutional Network (GCN)^27^, a DL method explicitly incorporating brain topology. These models were chosen to represent the spectrum of existing strategies commonly applied to classify outcomes, i.e., responders vs. nonresponders derived from ΔEM in our case. This evaluation was performed on ACT and CogTE, respectively; sample characteristics and protocol details are presented in **Table S1**. All models were trained using rsfMRI collected at T1 and T2. For non-FM methods (SVM, MLP, and GCN), FC features were derived separately at each timepoint and concatenated to encode longitudinal information. This design follows standard practice in conventional rsfMRI modeling^28,29^, where FC provides a stable, low-dimensional representation suitable for vector- or graph-based classifiers that do not operate directly on raw time-series data. The rsfMRI FMs used ROI-level time-series as input, which were temporally segmented into fixed-length patches and embedded as tokens (see **Methods**).

**Figure 2a** and **Table S2** summarize the model performance across benchmark comparisons. BrainLM outperformed other models, particularly on the CogTE dataset (*ACC: 80.3% ± 1%, F1: 88.2% ± 1%*). The performance advantage of BrainLM was also demonstrated on the multi-site ACT dataset (*ACC 65.2% ± 5%, F1 69.1% ± 5%*) compared to other models. In contrast, BrainJEPA did not achieve good performances across studies (*CogTE: ACC 60.4% ± 11%, F1 64.1% ± 7%; ACT: ACC 50.8% ± 13%, F1 53.4% ± 8%*). One possible explanation for this performance difference lies in their respective representation learning objectives: BrainJEPA’s cross-view consistency constraint may be less suited to the high variability characteristic of older adults’ BOLD signals, though this hypothesis warrants direct empirical validation through ablation or probing analyses; whereas, BrainLM’s autoregressive sequence modeling objective accommodates heterogeneous temporal patterns, making it more robust to modeling the neural dynamics among older adults.

**Figure 2.**
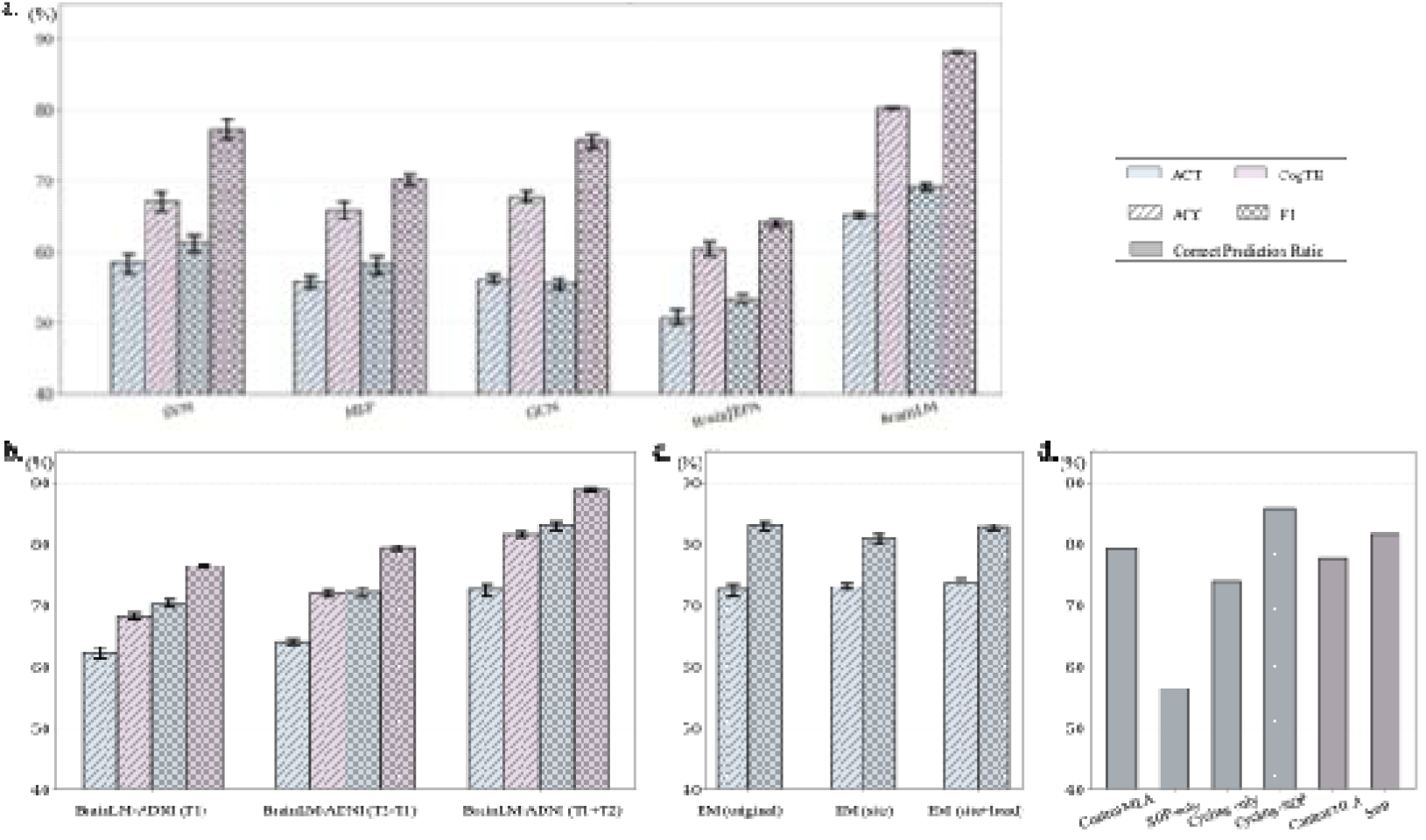
Evaluation of rsfMRI FMs for classifying intervention-related episodic memory response. **(a)** Benchmark comparison of classification performance between rsfMRI FMs (BrainLM, BrainJEPA) and conventional ML and DL approaches (SVM, MLP, GCN) across two independent intervention cohorts (CogTE and ACT). Models are trained to classify participants as episodic memory responders vs. non-responders, based on pre-post changes in EM, using baseline (T1) and post-intervention (T2) rsfMRI data. ACC and F1-score are reported as mean ± standard deviation across stratified cross-validation folds. **(b)** Performance of the BrainLM-ADNI under different timepoint combinations. Classification results for *T1 + T2, T1-only*, and *T2 − T1* input settings are shown for CogTE and ACT datasets (mean ± standard deviation across cross-validation folds). **(c)** Robustness of the adapted FM to common neuroimaging confounders in the multi-site ACT dataset. Classification performance of BrainLM-ADNI is shown for original data, data after site harmonization, and data after both site harmonization and head-motion regression. **(d)** Classification performance of BrainLM-ADNI stratified by intervention arm in both CogTE and ACT studies. Chi-square tests indicate no significant association between intervention assignment and classification correctness. **NOTE:** rsfMRI, resting-state functional magnetic resonance imaging; FM, foundation model; ML, machine learning; DL, deep learning; SVM, support vector machine; MLP, multilayer perceptron; GCN, graph convolutional network; ACT, Aerobic exercise and Cognitive Training study; CogTE, Cognitive Training and Exercise study; ADNI, Alzheimer’s Disease Neuroimaging Initiative; ACC, accuracy.

### Domain adaptive fine-tuning on ADNI dataset

To address Q2, we examined whether clinically informed adaptation and joint modeling of baseline and post-intervention rsfMRI improve intervention-relevant representations. Specifically, the best-performing BrainLM was fine-tuned using a supervised classification task distinguishing group with Alzheimer’s disease (AD) from healthy control (HC), named as BrainLM-ADNI. The design was motivated by the assumption that intervention-related EM improvement reflects, at least in part, a reversal or modulation of neural patterns of EM difference between AD and HC, making the HC-to-AD contrast a biologically meaningful axis for adaptation. Moreover, the differences between AD and HC are more neurobiologically distinct and stable than those between preclinical AD, such as MCI, offering a stronger supervisory signal for domain adaptation.

The resulting BrainLM-ADNI model was then evaluated on the ACT and CogTE datasets using the same downstream inference protocol as in **Section 2.1**. As shown in **Figure 2b**, BrainLM-ADNI demonstrated a substantial performance boost, particularly on the more challenging ACT dataset (*CogTE: ACC 81.6% ± 5% [*↑*1.3%], F1 88.8% ± 3% [*↑*0.6%]*); *ACT: ACC 72.6% ± 13% [*↑*7.4%], F1 83.1% ± 10% [*↑*14%]*), which suggests that the knowledge about aging-related brain patterns learned from ADNI was successfully transferred, equipping the model with a more refined understanding of the neural basis and its response to intervention, supporting our hypothesis. The consistent improvement across both datasets provides evidence that a clinically relevant, supervised fine-tuning step may still be critical for existing rsfMRI FMs when applying to a specialized clinical study.

To further characterize how the ADNI-adapted BrainLM leverages longitudinal information, we additionally evaluated model performance under different combinations of timepoints. Specifically, we compared the default “*T1+T2*” setting with a “*T1-only*” setting, in which downstream classification was based solely on baseline rsfMRI embeddings, and a “*T2–T1*” setting, in which residuals between post-intervention and baseline rsfMRI embeddings were used as input. As shown in **Figure 2b**, across two datasets, the default “*T1+T2*” configuration achieved the strongest and most consistent performance (*CogTE: ACC 81.6% ± 5%, F1 88.8% ± 3%, ACT: ACC 72.6% ± 13%, F1 83.0% ± 10%;*), indicating that jointly modeling baseline and post-intervention brain states provides complementary information beyond either baseline (*CogTE: ACC 68.3% ± 6%, F1 76.5% ± 5%, ACT: ACC 62.3% ± 11%, F1 70.4% ± 9%*,) or isolated change signals (*CogTE: ACC 72.0% ± 6%, F1 79.3% ± 5%, ACT: ACC 64.2% ± 5%*,, *F1 72.2% ± 9%*,) alone.

### Robustness to confounders concerning in neuroimaging and clinical trial studies

To address Q3, we evaluated whether the adapted rsfMRI foundation model remains robust to common sources of heterogeneity in intervention studies, including site variability, head motion, and intervention arm. Using the multi-site ACT dataset, we evaluated the performance of BrainLM-ADNI under three preprocessing conditions: (1) original data, (2) data after site harmonization, and (3) data after both site harmonization and regression of head motion parameters. As shown in **Figure 2c**, explicit correction for these confounders resulted in only modest differences in classification performance (*original: ACC 72.6% ± 13%, F1 83.0% ± 10%; site-harmonized: ACC 73.2% ± 7%, F1 81.0% ± 8%; site + head-motion corrected: ACC 73.8% ± 8%, F1 82.7% ± 6%*). Overall, these results indicate that explicit confound correction yields only modest performance changes, consistent with robustness to site-related heterogeneity and motion artifacts.

Additionally, we examined whether classification performance differed across intervention arms (see **Figure 2d**) in both studies. We performed a Chi-square test on intervention group assignment and classification performance (i.e., whether being correctly classified or not) for each participant. The classification performance displayed no significant difference across intervention arms for either study (*CogTE: χ^*2*^=0.01, p=0.92; ACT: χ^*2*^=5.44, p=0.14*). These results suggest that BrainLM-ADNI maintains comparable classification performance across intervention arms, further supporting its robustness.

### Identifying robust latent, multivariate neural patterns

To address Q4, we tested whether FM-derived embeddings support identification of latent neural patterns that are structured consistently across independent intervention studies. Specifically, we first applied Partial Least Squares (PLS) to characterize the relationship between brain activity (via amplitude of low frequency fluctuations; ALFF maps) and FM embeddings derived from the BrainLM-ADNI at T1 and T2, respectively, as well as components that differentiated responders from non-responders. This analysis was performed independently for both the ACT and CogTE studies. PSL components were selected based on their strong embedding-brain coupling and ability to differentiate responders from non-responders, providing a representative view of the embedding–brain relationship. Across both ACT and CogTE, for each timepoint (T1 and T2), the corresponding PLS component consistently demonstrated significant associations between FM-derived embeddings and ALFF-derived brain activity (**Figure 3a-b**). Notably, components identified at each timepoint independently contributed to this differentiation, indicating that relevant information is distributed across timepoints.

**Figure 3.**
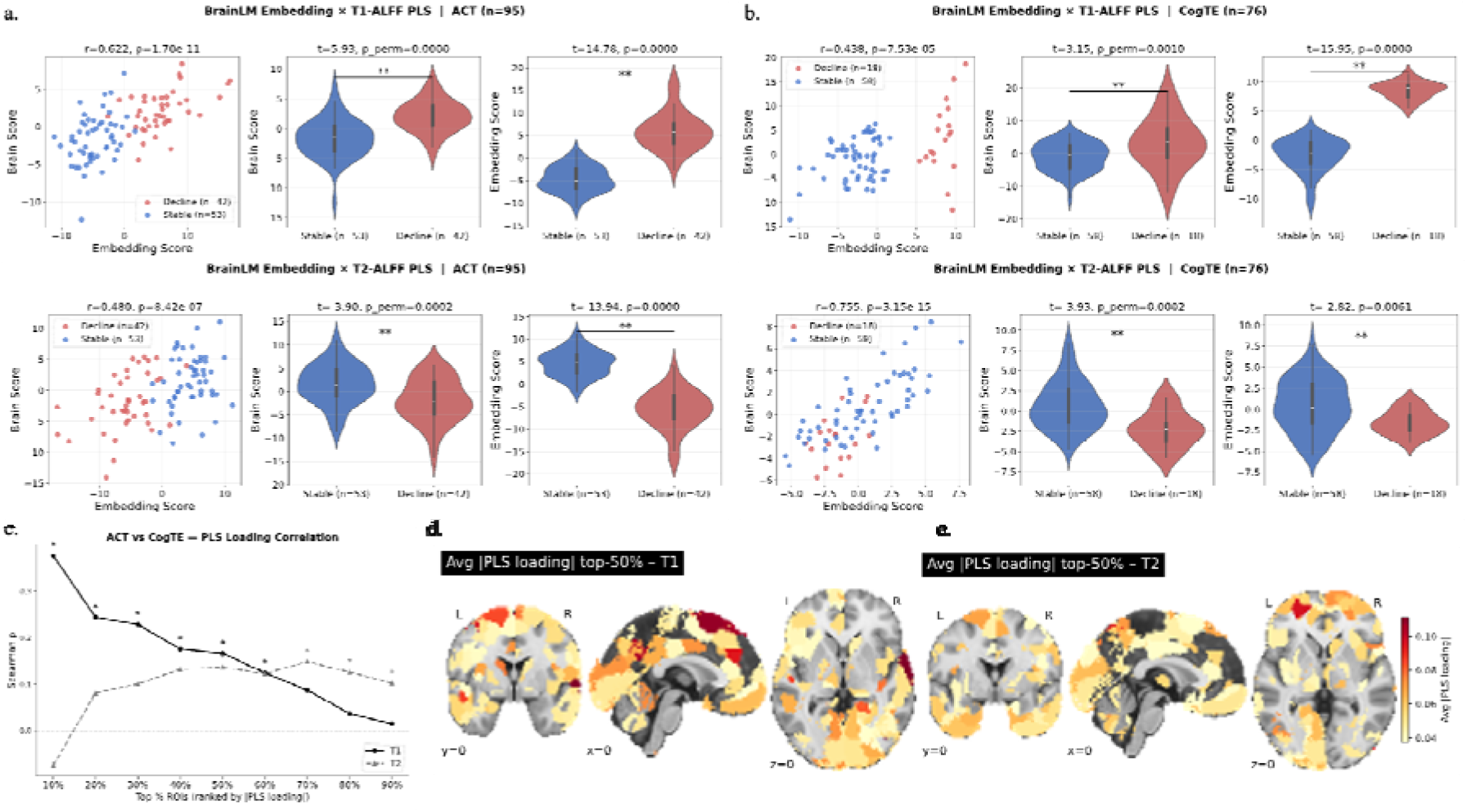
PLS-derived embedding-brain associations and cross-study spatial patterns. (a-b) ACT (a) and CogTE (b) PLS results. Top: T1; bottom: T2. Left: relationship between embedding scores and brain scores. Middle: group differences in brain scores. Right: group differences in embedding scores. Points are colored by responder status (stable vs. decline). Reported statistics correspond to two-sample t-tests on subject-level PLS scores. For each study and timepoint, the component shown represents the dominant embedding-brain association capturing group differentiation. (c) Cross-study correlations of PLS brain loading patterns across increasing thresholds of top-weighted regions (ranked by absolute loading). (d-e) Cross-study average spatial distribution of top-weighted PLS loadings at baseline (T1; left) and post-intervention (T2; right), shown in standard anatomical views and illustrating shared spatial organization of latent neural patterns across the two intervention studies.

Next, we examined cross-study consistency of PLS-derived brain loading patterns between ACT and CogTE (**Figure 3c**). At T1, similarity was highest among the top-ranked regions (peaking at top-10%) and gradually decreased as more regions were included, indicating that correspondence was driven by a focal set of dominant regions. At T2, similarity was minimal at the top ranks but peaked at intermediate thresholds (around top-50%), suggesting a more distributed pattern across regions. This shift indicates that baseline correspondence is concentrated in a core set of regions, likely reflecting aging-related constraints shared across studies, whereas post-intervention correspondence is more distributed and context-dependent, consistent with heterogeneity in interventions and cohorts rather than a lack of generalizability. To further explore the spatial organization of these patterns, we examined the distribution of top-50% PLS loadings across studies (**Figure 3d–e**). At T1, the highest-weighted regions covered brain regions primarily across default mode network, visual network, and frontoparietal network, especially highlighted in precuneus, paracingulate, and medical prefrontal regions, which are key networks involved in cognitive decline in late-life^30,31^. At T2, loading patterns became more spatially diffuse, consistent with the more distributed cross-study correspondence observed post-intervention.

## Discussion

### Summary of key findings across research questions

The overarching goal of this work was to identify robust, latent multivariate neural patterns of neuroplastic potential that generalize across studies and explain individual differences in cognitive response. Our findings across four research questions converge to support this goal. First, rsfMRI FMs, particularly the ADNI-adapted BrainLM, consistently outperformed established ML approaches in predicting individual treatment response across two independent intervention studies, even in small and heterogeneous samples, indicating that FM-derived embeddings capture meaningful intervention-related signal beyond predefined features such as FC. This finding indicates that FM-derived embeddings capture meaningful intervention-related signal beyond noise and substantially beyond representations based on predefined features such as FC. Second, clinically informed domain adaptation via ADNI fine-tuning was essential, and jointly modeling baseline and post-intervention states provided complementary longitudinal information, suggesting that intervention effects are encoded at a latent spatiotemporal representational level rather than as localized or static changes. Third, these representations demonstrated robustness across studies with differing designs, populations, and intervention modalities, suggesting that the models capture generalizable neural patterns relevant to cognitive response rather than study-specific artifacts. Importantly, architectural choice and domain adaptation played a critical role, with ADNI fine-tuning improving performance and BrainLM outperforming BrainJEPA, highlighting the importance of aligning pretrained representations with aging-related neural variability. Finally, interpretable embedding-brain relationships revealed consistent differentiation between responders and non-responders with structured cross-study organization, providing evidence that rsfMRI FMs can reliably identify latent neural patterns relevant to cognitive intervention outcomes.

### Identifying robust latent neural patterns with rsfMRI FMs in local intervention studies

We emphasize a **three-condition framework for establishing robust latent, multivariate neural patterns** in cognitive aging intervention research: First, the analytical model must learn brain representations that preserve fine-grained spatiotemporal structure beyond predefined features, despite noise, heterogeneity and limited sample size; Second, these representations must demonstrate behavioral relevance by meaningfully relating to individual cognitive change following intervention; Third, the identified neural representations must exhibit consistency across independent studies, indicating shared neural signature rather than study-specific artifacts. Our findings satisfy the first condition. FM-derived embeddings captured subtle spatiotemporal patterns that are difficult to recover from predefined connectivity features or models trained from scratch on small intervention datasets, a critical advantage in aging populations where intervention effects are modest and embedded within large inter-individual variability. Importantly, large-scale pretraining alone was insufficient; ADNI-based fine-tuning acted as an essential intermediate step, aligning general representations with aging-related neural variance and enabling transfer to intervention settings.

The second condition was met through PLS-based latent component generation, which mapped FM-derived embeddings onto low-dimensional latent variables directly coupled to behavioral outcomes. Unlike fixed anatomical targets or predefined connectivity metrics, these latent variables allow intervention-related variation to emerge as distributed network-level patterns, which are well-suited to aging populations where neural changes are diffuse, modest, and not easily localized to canonical regions. Each selected PLS component demonstrated significant group separation between responders and non-responders, confirming behavioral relevance across studies and timepoints.

The third condition was supported by structured cross-study consistency in PLS loading patterns. At baseline, correspondence was concentrated in a focal set of highest-weighted regions anchored in posterior medial cortex, reflecting shared neurobiological constraints across aging cohorts. Post-intervention, correspondence emerged at broader thresholds, indicating that intervention-driven neural changes are more distributed and context-dependent. Together, these patterns define candidate neural signatures of intervention-relevant variation that can be examined and validated across independent studies. We further conceptualize these insights into a unified framework (**Figure 4**).

**Figure 4.**
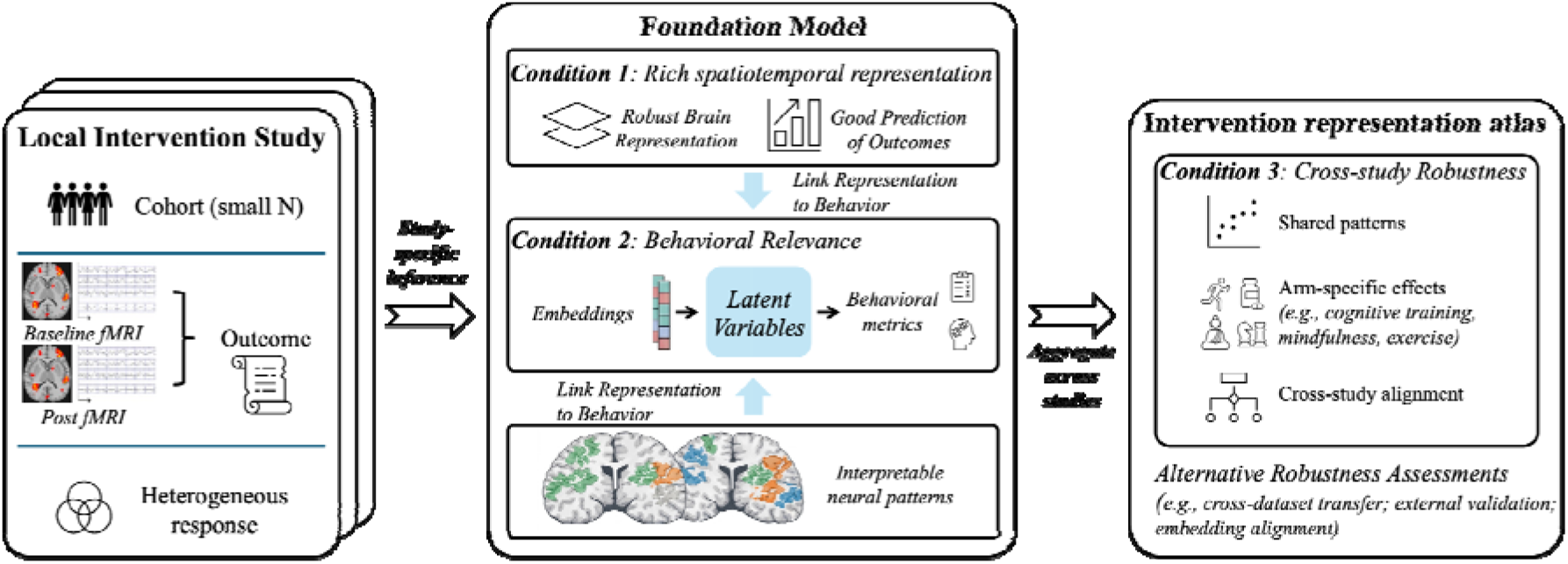
Three-condition framework for identifying robust and generalizable neural patterns in cognitive aging intervention studies. rsfMRI FMs are applied to local intervention studies characterized by small sample sizes and heterogeneous responses. Under Condition 1 (rich spatiotemporal representation), FMs extract latent neural embeddings from BOLD time series that capture distributed dynamics beyond predefined features, enabling robust representation learning and improved prediction of outcomes. Under Condition 2 (behavioral relevance), these embeddings are linked to behavioral metrics through multivariate analysis, yielding latent variables that characterize individual differences in intervention response. These latent representations further support the identification of interpretable neural patterns. Under Condition 3 (cross-study robustness), neural patterns are evaluated across independent studies to identify shared structures, distinguish arm-specific effects, and enable cross-study alignment. Together, these steps define an intervention representation atlas that captures both shared and study-specific neural signatures. Alternative approaches, including cross-dataset transfer, embedding alignment, and external validation, may provide complementary assessments of generalizability.

**Figure 5.**
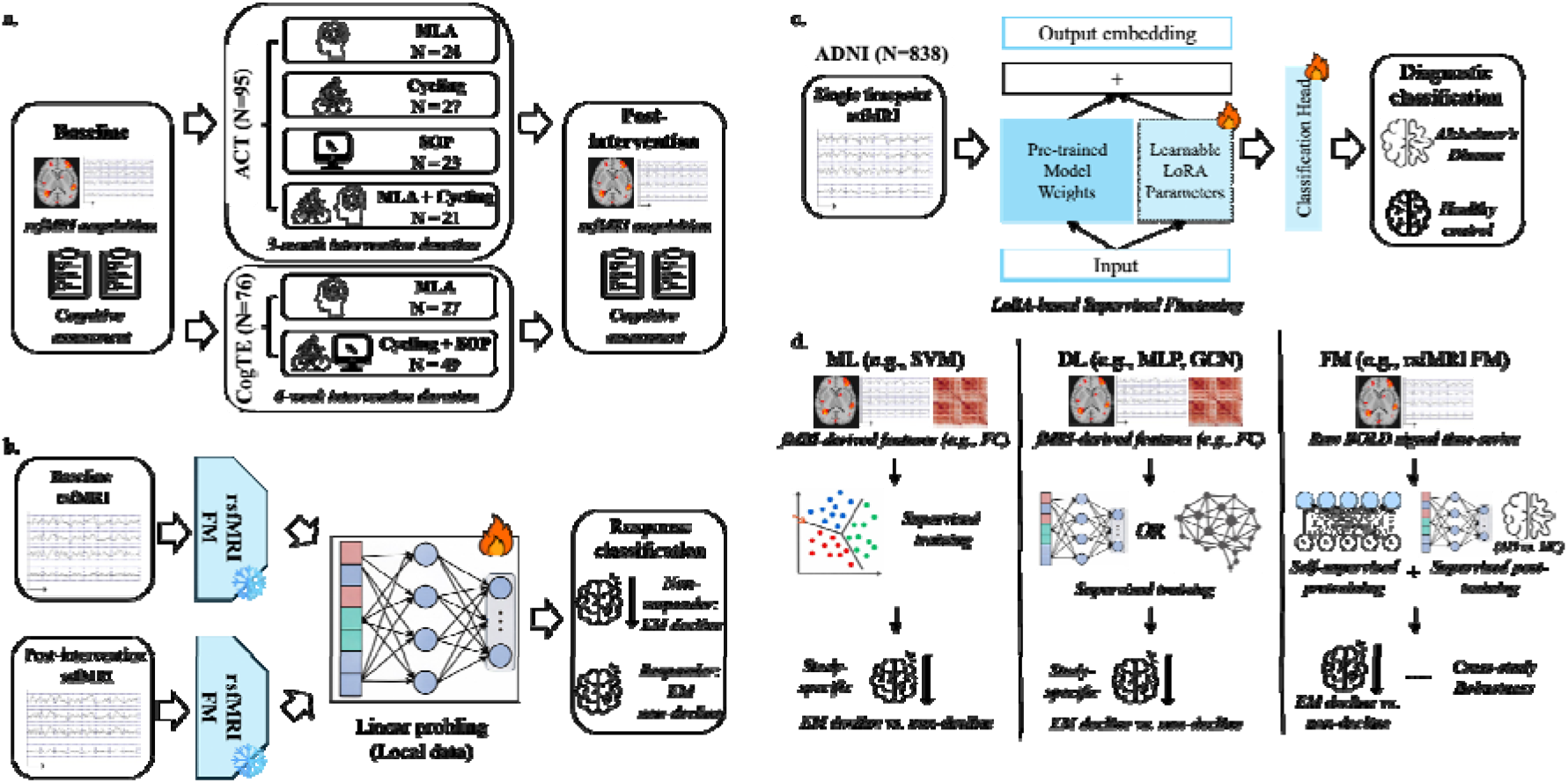
Study datasets and implementation of rsfMRI FMs and comparison methods. (a) The ACT and CogTE studies serve as independent intervention cohorts for downstream evaluation and assessment of model generalizability across intervention paradigms and study designs. (b) Domain-adaptive finetuning of BrainLM using the ADNI dataset via LoRA. Supervised finetuning is performed to incorporate aging- and pathology-related brain patterns, yielding the adapted BrainLM-ADNI model that is subsequently used as a frozen feature extractor in all downstream intervention analyses. (c) Linear probing strategy for deploying pretrained rsfMRI FMs in longitudinal intervention studies. The rsfMRI data collected at baseline (T1) and post-intervention (T2) are processed independently through a frozen FM with shared weights, and CLS-token embeddings from each timepoint are concatenated to explicitly encode longitudinal neural change for downstream inference of EM outcomes. (d) Architectures of conventional comparison models. FC features derived from rsfMRI data at T1 and T2 are used to train an SVM, an MLP, and a dual-stream GCN that explicitly models longitudinal brain network changes. NOTE: rsfMRI, resting-state functional magnetic resonance imaging; FM, foundation model; ACT, Aerobic exercise and Cognitive Training study; MLA, mental leisure activity; CogTE, Cognitive Training and Exercise study; ADNI, Alzheimer’s Disease Neuroimaging Initiative; EM, episodic memory; FC, functional connectivity; T1/T2, baseline/post-intervention timepoints; SVM, support vector machine; MLP, multilayer perceptron; GCN, graph convolutional network; LoRA, low-rank adaptation; AD, Alzheimer’s Disease; HC, healthy control.

Looking forward, an important next step is to aggregate such latent neural signatures across diverse intervention modalities (e.g., aerobic exercise, neuromodulation, mindfulness, and cognitive training) to construct a shared reference space of intervention-associated brain patterns. Within this space, FM-derived embeddings could serve as a common coordinate system for comparing individuals and interventions, enabling evaluation of how an individual’s baseline neural representation aligns with patterns associated with different response profiles. While this application is prospective, the framework outlines a path from post-hoc analyses to prescriptive medicine: a scalable, data-driven approach to personalizing interventions based on an individual’s latent neuroplastic potential, not on group-average effects.

### Practical considerations for identifying robust latent neural patterns with rsfMRI FMs

For cognitive aging intervention research to make cumulative progress toward mechanistic insight and personalized treatment, it is essential for the field to consistently identify and compare latent, multivariate neural patterns across studies, rather than relying on isolated within-trial findings. In this context, applying rsfMRI FMs in intervention studies is best understood as a practical strategy for satisfying the three conditions required for robust latent pattern identification, rather than as a single fixed pipeline. Under realistic constraints such as limited sample size, heterogeneous responses, and domain mismatch, each condition can be supported by both the approach used here and alternative solutions. For the first condition, we used pre-trained rsfMRI FMs operating on BOLD time series and further adapted them using ADNI to better align the representation space with aging-related neural variation. Alternative solutions could include other pretrained sequence, graph-based^32^, or multimodal models^33^, to preserve fine-grained neural dynamics beyond predefined summary features. For the second condition, we evaluated whether FM-derived embeddings predicted intervention response and used multivariate decomposition to identify latent components coupled to EM change. Alternative approaches could include other brain-behavior linkage methods, such as canonical correlation analysis or supervised latent variable models, to retain representations that explain meaningful individual cognitive variation, not just model performance. For the third condition, we assessed consistency across two independent intervention cohorts and examined robustness to confounders. Alternative approaches may include cross-dataset transfer, embedding alignment, or external validation, where identified patterns reflect shared neural structure rather than study-specific artifacts. In practice, these steps involve trade-offs among data access, model scale, interpretability, and engineering effort.

## Materials and Methods

### Overview of the analytical framework

To systematically evaluate whether rsfMRI foundation models can support robust and generalizable identification of intervention-relevant latent neural patterns in cognitive aging studies, we designed a multi-component framework (**Figure 1**). For each participant, rsfMRI data (both T1 and T2) were processed in a dual-stream architecture with shared model weights to extract latent embeddings for each timepoint. These embeddings were subsequently concatenated as input features to explicitly encode longitudinal neural change for downstream prediction of intervention response and subsequent identification of latent brain–behavior patterns. To ensure a fair and controlled evaluation, all downstream analyses were conducted within a stratified five-fold cross-validation framework that balanced outcome labels, intervention arms, and study sites when it is applicable. This design enabled direct comparison between FMs and conventional machine learning and deep learning approaches under identical data splits and evaluation protocols. Model performance was primarily assessed using Accuracy (ACC) and F1-score, reflecting overall correctness and the balance between precision and recall, respectively. To provide a complementary assessment of discriminative ability and class-specific performance, Area Under the ROC Curve (AUC), Sensitivity, and Specificity were also reported (see results in **Supplementary Materials Table S2**). Performance metrics were averaged across cross-validation folds, and the same evaluation protocol was applied consistently to all ML, DL, and rsfMRI FM approaches.

### Study cohorts and preprocessing

Three independent datasets were included in this work for model deployment and evaluation. The ADNI dataset was used exclusively for domain-adaptive fine-tuning of the FM. The ADNI dataset comprises older adults spanning the cognitive spectrum from cognitively normal, MCI, to AD. In this study, ADNI participants diagnosed with AD and cognitively healthy controls were selected to construct a supervised fine-tuning task that captures aging- and pathology-related brain patterns. No ADNI data were included in the downstream model evaluation. The ACT study is a multi-site randomized controlled trial involving older adults at risk for AD, with participants assigned to aerobic exercise, cognitive training, combined intervention, or active control arms over six months. Resting-state fMRI and episodic memory assessments were collected at T1 and T2. The CogTE study is a single-site randomized controlled trial with a six-week cognitive training intervention and an active control condition, following a similar pre-post assessment design. Both ACT and CogTE datasets target older adults with elevated risk for cognitive decline and serve as independent cohorts for evaluating model generalizability across intervention paradigms and study designs (see **Figure 3a**). Both datasets exhibited substantial inter-individual variability in EM change, with samples of responders and non-responders in each intervention arm (see **Supplementary Materials Table S1**).

MRI acquisition details for ADNI, ACT, and CogTE have been reported previously and are summarized in **Supplementary Materials Table S1**. All imaging data were preprocessed using the standardized *fMRIprep* pipeline^34^ (version 24.1.0), which is based on *Nipype* v1.8.6. For each participant, T1-weighted anatomical images underwent skull stripping, tissue segmentation, surface reconstruction, and spatial normalization to Montreal Neurological Institute (MNI) space. RsfMRI images were corrected for head motion and slice timing, co-registered to the anatomical image, and normalized to MNI space. Susceptibility distortion correction was applied for the ACT dataset using estimated field maps from two echo-planar imaging references with *TOPUP*^35^ that were available only for this study. Following *fMRIPrep*, additional denoising was performed using *Nilearn* (version 0.11.0). This included regression of twelve head motion parameters, five principal components derived from white matter and cerebrospinal fluid signals using the *aCompCor* approach, and temporal band-pass filtering (0.01–0.1 Hz). Finally, spatial smoothing was applied using a 6-mm full-width-at-half-maximum Gaussian kernel. Identical preprocessing procedures were applied across all datasets to minimize confounding differences arising from data handling and to ensure consistency in downstream model evaluation.

### Resting-state fMRI foundation model deployment

**BrainLM** is designed to model whole-brain resting-state activity as an inference sequence, analogous to autoregressive language modeling. During pretraining, BrainLM learns to predict future spatiotemporal patterns of brain activity conditioned on past observations, enabling it to capture long-range temporal dependencies and distributed neural dynamics. BrainLM operates directly on preprocessed rsfMRI time-series data, where voxel- or parcel-wise BOLD signals are temporally segmented into fixed-length patches of 20 datapoints and embedded as input tokens to the transformer encoder. Although three pretrained variants (13-/111-/650-million weights) were described in the original paper, only the 13-million weight model was publicly released and was therefore used in this study.

**BrainJEPA** follows a joint embedding inference architecture. Its primary objective is to learn shared latent representations by enforcing consistency between embeddings derived from neighboring spatiotemporal views of the brain. BrainJEPA is trained to align representations of partially observed brain states, encouraging invariance across local temporal or spatial perturbations. As a result, it emphasizes representation consistency and local coherence rather than explicit temporal forecasting. BrainJEPA takes preprocessed rsfMRI time-series as input, formatting the signals into sequences of 160 time points and segmenting them into non-overlapping patches of size 16. Although three pretrained variants (22-/86-/307-million weights) were described in the original paper, only the 22-million weight model was publicly released and was therefore used in this study.

To deploy these rsfMRI FMs to local intervention studies without retraining the backbone, a linear probing strategy was employed. For each participant, rsfMRI data from T1 and T2 were independently passed through the frozen FM using shared weights. From each session, the final hidden representation corresponding to the Classification (CLS) token was extracted as a fixed-dimensional embedding. These embeddings summarize the participant’s brain state at each timepoint in the learned representation space. The T1 and T2 embeddings were then concatenated to form a longitudinal feature vector explicitly encoding pre-post neural change. This fused representation was used as input to a lightweight, task-specific classifier trained to ΔEM. During linear probing, all FM weights remained frozen, and only the weights of the downstream classifier were optimized. This setup isolates the representational capacity of the pretrained rsfMRI FMs and allows a controlled comparison of their ability to extract longitudinal representations that support both intervention-response prediction and downstream discovery of latent brain-behavior patterns in small-sample intervention datasets (**Figure 3b**). Detailed training configurations are described in **Supplementary Materials**.

### Model fine-tuning

To adapt the pretrained rsfMRI FM to aging-related neural variation before downstream intervention analysis, supervised fine-tuning was performed on the ADNI dataset as a clinically informed representation adaptation step. This fine-tuning step was applied exclusively to BrainLM, based on its superior transfer performance in zero-shot and linear probing settings. Fine-tuning was formulated as a binary classification task distinguishing participants diagnosed with AD vs. HC. This task was selected to introduce aging- and pathology-relevant supervisory signals into the pretrained representation space, without altering the downstream intervention datasets. Only ADNI data were used during fine-tuning, and no ADNI subjects were included in any downstream evaluation. To reduce overfitting risk and computational cost, lightweight Low-Rank Adaptation (LoRA) modules were inserted into the query, key, value, and output projection layers of the self-attention blocks in the BrainLM transformer. Most pretrained FM weights remained frozen, while the final transformer block was unfrozen to allow limited adaptation of higher-level representations. It enables the model to incorporate domain-specific information while preserving the general spatiotemporal representations learned during large-scale pretraining. During fine-tuning, the downstream classifier consisted of a two-layer MLP operating on the CLS token embedding produced by the BrainLM. Model optimization was conducted using the AdamW optimizer with differential learning rates applied to the adapted rsfMRI FM weights and the downstream classifier. A cosine learning rate scheduler with warmup was used to stabilize training. Early stopping was applied based on validation loss to prevent overfitting, and the model checkpoint with the lowest validation loss was selected. After fine-tuning, the adapted BrainLM model (denoted as BrainLM-ADNI) was used as a frozen feature extractor for all downstream intervention-response prediction and latent pattern analyses. No further weight updates were performed during linear probing on the ACT and CogTE datasets (see **Figure 3c**).

### Compared ML and DL methods

Support Vector Machine (SVM) served as a strong traditional machine learning method. For each participant, FC matrices were computed separately for T1 and T2 rsfMRI scans. The upper triangular elements of each FC matrix were flattened into feature vectors and concatenated across timepoints to explicitly encode longitudinal change. The resulting feature vector was used as input to a linear SVM classifier (see **Figure 3d**). A linear kernel was chosen to reduce overfitting risk under limited sample sizes. The regularization parameter *C* was optimized via grid search within the training folds of the cross-validation procedure.

Multilayer Perceptron (MLP) represents a vector-based deep learning method capable of modeling nonlinear relationships. The input features were the same as those used for the SVM, consisting of concatenated upper-triangular FC vectors from T1 and T2. The MLP architecture comprised three fully connected layers with ReLU activation functions and dropout for regularization (see **Figure 3d**). Hyperparameters, including hidden layer dimensions, dropout rates, and learning rate, were tuned within the training folds to mitigate overfitting.

Graph Convolutional Network (GCN) was implemented to explicitly incorporate brain network topology and model longitudinal graph-structured data. Brain regions of interest (ROIs) were treated as nodes in a graph, with edges defined by a shared adjacency matrix constructed from a control-group functional connectivity template. For each participant, node features were derived from FC matrices at T1 and T2, resulting in two graph inputs representing baseline and post-intervention brain states. A dual-stream GCN architecture with shared weights was used to process the T1 and T2 graphs in parallel. Each stream consisted of stacked graph convolutional layers followed by graph-level pooling to produce timepoint-specific embeddings. These embeddings were then fused by concatenation along with their difference to explicitly encode longitudinal change. The fused representation was passed to a fully connected output layer for classification (see **Figure 3d**). Model parameters were optimized using the Adam optimizer, and training was conducted within the same stratified cross-validation framework as all other models.

### Post-hoc latent pattern and cross-study consistency analyses

Partial Least Squares (PLS) was performed to identify multivariate associations between FM-derived embeddings and whole-brain neural activity in each dataset independently. For each time point (T1, T2), an embedding matrix *X* ∈ ℝ^*N×d*^(*d* = 128) was constructed from the penultimate layer of the downstream classifier, and a brain matrix *Y* ∈ ℝ ^*N×p*^(*p* = 424) was derived from parcel-wise ALFF values based on the A424 atlas. All variables were z-scored prior to analysis. Canonical PLS (n_components = 3, max_iter = 2000) decomposed the cross-covariance *X*^T^*Y* into latent variables, yielding embedding scores (*u* = *Xw*_*x*_) and brain scores (*v* = *Xw*_*y*_), where *w*_*x*_ and *w*_*y*_ denote the respective weight vectors. The brain loading vector w_*y*_ was used to quantifies each ROI’s contribution for subsequent spatial interpretation. The primary component was defined as the one with the largest absolute between-group t-statistic on outcomes (responder vs. non-responder). This selection strategy prioritizes latent dimensions most relevant to individual intervention response. Group differences in brain and embedding scores were assessed using independent-samples t-tests. To account for potential violations of parametric assumptions, a non-parametric permutation test (5,000 permutations) was conducted for the brain score of the primary component, with empirical p-values computed from the proportion of permuted | *t* | exceeding the observed value. Effect sizes were reported as Cohen’s d. To evaluate the consistency across ACT and CogTE trials, we also tested correlation of loadings between studies within each timepoints. The ROI regions were ranked by absolute values of PLS loadings within studies and timepoints. We selected the union of ROIs of two trials selected by ranking percentage from 10% to 90%, with 10% as interval. Based on these sets of ROIs, we tested between-study correlation at each percentage thresholds.

### Data, Materials, and Software Availability

Data from the ADNI are publicly available and can be accessed through the ADNI data-sharing platform following approval of a data use application. Data from the ACT and CogTE intervention studies contain information that could compromise participant privacy and are therefore not publicly available; de-identified data may be made available from the corresponding authors upon reasonable request and subject to institutional approval. All imaging data were processed using publicly available software, including *fMRIPrep* and *Nilearn*. Publicly released checkpoints of BrainLM and BrainJEPA were used in this study. Code supporting the analyses and results reported here will be made publicly available upon publication.

## Acknowledgments

The project was supported by NIH NR015452 and Stanford HAI grant.

## Notes

**Competing Interest Statement:** Authors claimed no conflict of interest.

### Competing Interest Statement

The authors have declared no competing interest.

### Summary of Updates

The revised manuscript has been extensively reworked in several important ways. First, we reframed the study. It is now a framework for discovering robust and generalizable latent neural targets of intervention response in cognitive aging, rather than primarily a model-evaluation paper. Second, we clarified the methodological logic. We explicitly positioned model fine-tuning as a clinically informed representation adaptation step that comes before downstream intervention-response prediction and latent pattern discovery. Third, we substantially revised and expanded the post hoc analyses. The new analyses better characterize latent brain-behavior structure and cross-study consistency. These now include multivariate decomposition of foundation model embeddings and their relationship to episodic memory change across two independent intervention trials. Fourth, we formalized a three-condition framework. This includes rich spatiotemporal representation, behavioral relevance, and cross-study robustness for clinically meaningful latent target discovery. This framework now guides the revised title, abstract, introduction, discussion, and figures.

