## Supporting materials for "Resting-state fMRI foundation models enable robust and generalizable latent neural target discovery in cognitive aging interventions"

#### 1. Sample characteristics and protocol details of CogTE and ACT dataset

|  | CogTE (N=76) | | ACT (N=95) | | | |
| --- | --- | --- | --- | --- | --- | --- |
|  | SOP (n=49) | MLA (n=27) | ACT(n=21) | SOP (n=23) | Cycling (n=27) | MLA (n=24) |
| EM at T1 | 38.71±13.67 | 39.11±12.43 | 43.10±11.53 | 44.61±9.67 | 39.37±12.45 | 42.13±13.33 |
| EM at T2 | 41.76±14.64 | 43.85±12.16 | 43.76±13.43 | 41.91±12.85 | 39.37±11.48 | 41.33±10.36 |
| ΔEM  (T2-T1) | 3.04 ±10.38 | 4.74±12.23 | 0.67±10.46 | -2.70±9.45 | 0.00±10.52 | -0.79±11.26 |
| Number of responders (n, %) | 39, 53% | 19, 26% | 13, 14% | 11, 12% | 17, 18% | 12, 13% |
| Intervention duration | 6-week | | 28-week | | | |
| Intervention protocol | Computerized visual processing speed training (VSOP) which emphasized on processing speed and attention | Leisure activities including online word search, Sudoku, and Free-Cell, a variation of solitaire | Participants followed the cycling protocol first, immediately followed by the SOP protocol within the same session. Each ACT session lasted 80 minutes, combining the durations of cycling-only and SOP-only protocols. | Five attention- and processing-speed-focused games (Eye for Detail, Hawk Eye, Visual Sweeps, Double Decision, and Target Tracker). SOP session durations decreased from 50 minutes to 30 minutes over time. | Moderate-to-vigorous intensity cycling using a non-linear progression approach, with intensity ranging from 50–75% of heart rate (HR) reserve (based on exercise testing during screening) or 11–15 on the Rating of Perceived Exertion (RPE) scale. Sessions lasted 30–50 minutes. | Stretching and mental leisure activities for a total of 80 minutes per session. Stretching included seated movements and static stretches matching the duration of cycling-only sessions. Mental leisure activities such as word search, Sudoku, and solitaire games, matched the duration of SOP-only sessions. |
| MRI acquisition details | MRI data were collected under a 3T Siemens TrioTim scanner. An echo-planar imaging (EPI) MPRAGE sequence was conducted to collect high-resolution structural anatomical image: repetition time/echo time (TR/TE) = 2350/3.44ms, TI=1100ms, FA=7, acquisition matrix 256*256mm, slice thickness = 1mm, resolution = 1 mm isotropic, 192 slices. A gradient echo-planar imaging sequence was conducted to collect resting-state functional images: TR/TE = 2500/30ms, FA=90, acquisition matrix 64*64mm, slice thickness = 4 mm, 37 axial slices. The duration of rs-fMRI scan was 5 minutes, with an eyes-open protocol. | | MRI data were collected under Siemens 3T Prisma (VE11C) with a 64-channel head coil. A single-shot echo-planar imaging (EPI) MPRAGE sequence was conducted to collect high-resolution structural anatomical image: repetition time/echo time (TR/TE) = 1400/2.34 ms, slice thickness = 1 mm, resolution = 1 mm isotropic, 192 slices, PE acceleration = GRAPPA, flip angle = 8°, orientation = sagittal, echo spacing = 7 ms, field of view (FOV) = 256 mm. A five-minute gradient EPI multiband (MB) sequence was conducted to collect resting-state functional images: TR/TE = 1010/44 ms, slice thickness = 2 mm, resolution = 2 mm isotropic, R = 1, MB acceleration factor = 8, flip angle = 70°, orientation = transversal, echo spacing = 0.56 ms, FOV = 256 mm, acquisition matrix = 128 x 128, 80 slices, 253 volumes, acceleration factor = 8. Slice acquisition order was interleaved. Participants were instructed to keep their eyes open. Field maps with opposite phase encoding directions were acquired for distortion correction. The same MRI acquisition protocol was applied across all three sites. | | | |

***Table S1.*** *Descriptive information of EM outcome and protocol details for CogTE and ACT study. T1, baseline; T2, post-intervention; SOP, Speed of Processing; ACT, Speed of Processing plus Cycling training; MLA, Mental Leisure Activity.*

#### 2. Performance results across benchmark comparisons.

| **Dataset** | **Method** | **ACC** | **F1** | **AUC** | **Sensitivity** | **Specificity** |
| --- | --- | --- | --- | --- | --- | --- |
| **ACT** | SVM | 58.4% ± 17% | 61.2% ± 15% | 48.6% ± 10% | 60.5% ± 12% | 46.8% ± 18% |
|  | MLP | 55.8% ± 11% | 58.2% ± 16% | 48.9% ± 11% | 63.8% ± 12% | 45.2% ± 17% |
|  | GCN | 56.2% ± 8% | 55.5% ± 10% | 52.2% ± 8% | 61.0% ± 10% | 48.5% ± 11% |
|  | BrainJEPA | 50.8% ± 13% | 53.4% ± 8% | 46.0% ± 12% | 58.5% ± 11% | 42.0% ± 16% |
|  | BrainLM | 65.2% ± 5% | 69.1% ± 5% | **61.9% ± 5%** | 75.0% ± 7% | 56.0% ± 9% |
|  | **BrainLM-ADNI** | **72.6% ± 13%** | **83.1% ± 10%** | 60.8% ± 14% | **76.1% ± 6%** | **59.0% ± 12%** |
| **CogTE** | SVM | 67.1% ± 17% | 77.2% ± 18% | 46.0% ± 6% | 62.5% ± 14% | 40.0% ± 19% |
|  | MLP | 65.8% ± 14% | 70.2% ± 10% | 48.7% ± 11% | 66.8% ± 11% | 44.0% ± 14% |
|  | GCN | 67.8% ± 10% | 75.6% ± 12% | 49.8% ± 12% | 65.0% ± 12% | 43.5% ± 15% |
|  | BrainJEPA | 60.4% ± 11% | 64.1% ± 7% | 44.2% ± 12% | 65.5% ± 12% | 41.0% ± 13% |
|  | BrainLM | 80.3% ± 1% | 88.2% ± 1% | 57.9% ± 8% | 78.0% ± 10% | 50.5% ± 6% |
|  | **BrainLM-ADNI** | **81.6% ± 5%** | **88.8% ± 3%** | **59.8% ± 11%** | **80.8% ± 8%** | **54.5% ± 10%** |

***Table S2.*** *Performance of different classification models on the ACT and CogTE intervention datasets. Values represent mean ± standard deviation across stratified cross-validation folds. Accuracy (ACC), F1-score, area under the curve (AUC), Sensitivity, and Specificity are reported. All models were evaluated using resting-state fMRI data acquired at baseline (T1) and post-intervention (T2) under identical training and evaluation protocols. Bold values indicate the optimal result.*

#### 3. Model training configurations

All model training and evaluation were conducted on the Google Cloud Platform using NVIDIA A100 GPUs with 40 GB memory. Deep learning models were implemented using PyTorch and PyTorch Geometric, while conventional machine learning baselines utilized Scikit-learn. Training was accelerated using CUDA Automatic Mixed Precision.

Two representational learning models were evaluated using publicly available checkpoints: (1) BrainLM, the 13-million parameter version was used; the model processes BOLD time-series segmented into fixed-length patches of 20 datapoints; (2) BrainJEPA, The 22-million parameter version was applied; the model processes sequences of 160 time points, segmented into non-overlapping patches of size 16.

For the downstream classification on the ACT and COGTE datasets, a linear probing protocol was employed. The classification head, a two-layer MLP, was trained for 30 epochs using the *Adam* optimizer, a learning rate of 1e-4, and the *BCEWithLogitsLoss* function with a batch size of 8. The input to this head was the concatenated *CLS* token embeddings from the T1 and T2 timepoints, which were extracted from a shared and frozen ADNI-tuned BrainLM backbone.

For ADNI finetuning, the default model consisted of a 4-layer, 4-head BrainLM *transformer* with a hidden size of 512. *LoRA* was employed with a rank of 32, alpha of 64, and dropout of 0.1, targeting the query, key, value, and dense layers of the attention mechanism. In addition to the *LoRA* modules, the final embedding layer of the transformer was also unfrozen for training. The classification head consisted of a two-layer *MLP* (512→256→2) with *LayerNorm*, *GELU* activation, and *dropout* (layer 1: p=0.3: layer 2: p=0.2). The model was trained for 500 epochs with a batch size of 16, using an *AdamW* optimizer and a weight decay of 0.01. A differential learning rate was used: 1e-5 for the backbone and *LoRA* parameters, and 1e-4 for the classification head. A cosine learning rate scheduler with a 10% warmup phase was applied. The checkpoint with the lowest validation loss was saved as the best model for the subsequent stage.
